# HIRA-SETDB1-H3K9me3 axis regulate chromatin architecture in chronic myeloid leukemia cells

**DOI:** 10.1101/2024.11.01.621462

**Authors:** Mayur Balkrishna Shirude, Anjali Devarajan, Debasree Dutta

## Abstract

Histone cell cycle regulator A (HIRA) confers chromatin accessibility and regulates developmental hematopoiesis. But whether HIRA displays similar role in leukemia, a condition caused by abnormalities during hematopoiesis, remain elusive. Here we show that HIRA interact with heterochromatin cluster in Chronic Myeloid Leukemia, K562, cells. FRAP, FLIM-FRET and ATAC-sequencing analysis revealed increased chromatin compaction, altered spatial distribution of chromatin towards nuclear periphery and loss in chromatin accessibility at the promoter and gene bodies upon downregulation of HIRA in K562 cells. Enhanced chromatin compaction was attributed to increased histone H3K9me3 level mediated by histone methyltransferase SETDB1. Incorporation of histone H3.3 within the *SETDB1* promoter in *HIRA*-knockdown cells induced SETDB1 expression. HIRA-SETDB1-H3K9me3 axis regulate the chromatin architecture in CML cells leading to inhibition of proliferation while induction in differentiation. We anticipate that the exploration of this axis would introduce new paradigm in understanding and targeting molecules that could influence CML disease progression.

## Introduction

Chromatin architectural changes are very dynamic and crucial for the cell fate decisions during cellular differentiation. Embryonic and hematopoietic stem cells (HSCs) differentiation process is accompanied by dramatic global chromatin architectural changes (Dixon et al. 2015). Differentiation of HSCs to blood cells show progressive chromatin condensation mediated by H3K9 methylation that gets redistributed towards nuclear periphery (Ugarte et al. 2015). Recent report from our lab demonstrated the role of histone chaperone Histone cell cycle regulator A (HIRA) in regulation of RUNX1 indispensable for hemogenic to hematopoietic transition (Majumder et al. 2015). HIRA, a replication independent histone chaperone associates with UBN1, CABIN1 and transiently with ASF1A to form a complex that incorporates H3.3 and marks active transcription (Ray-gallet et al. 2002; Pchelintsev et al. 2013; Tagami et al. 2004). H3.3 exhibits apparently contradicting roles at different genetic loci (Choi et al., 2024). HIRA has been shown to regulate chromatin accessibility and modulate the process of developmental hematopoiesis (Chen et al., 2020). But, whether HIRA could play a similar role in the chromatin accessibility during abnormal hematopoiesis, or leukemia, remain unexplored. Leukemia is a cancer of the blood and bone marrow. Different types of leukemia are being reported depending on the type of blood cell getting affected. Our earlier study demonstrated an induced expression of HIRA in chronic myeloid leukemia (CML) patient samples and in cell line derived from CML patients (Majumder et al. 2019). CML is a gradually advancing hematological and bone marrow disorder that often manifests at or after middle age and is exceedingly uncommon in children under the age of 15. The presence of fusion protein arising out of chromosomal translocation of t(9;22)(q34;q11.2), Breakpoint Cluster Region protein-Abelson Tyrosine-Protein Kinase 1 (BCR-ABL1) oncoprotein, results in constitutive defective tyrosine kinase activity that influences cell proliferation, apoptosis and survival among others. Starting with an asymptomatic chronic phase, it goes through accelerated phase and finally reaches the blast phase. Aberrant or uncontrollable proliferation and impaired differentiation marks leukemia and our earlier study demonstrated the role of HIRA in regulation of proliferation vs differentiation of CML cells (Majumder et al. 2019). Loss in HIRA level resulted in inhibition of proliferation while induction of differentiation to megakaryocyte fate (Majumder et al. 2019). Although a significant number of studies have been conducted in understanding the role of HIRA in developmental hematopoiesis, but, its role in leukemia have not been explored further. Chromatin architecture aberrations are linked to various human diseases, including cancer (Wang 2023). The induced expression of HIRA in CML patient samples and in cell line and its regulatory role in chromatin states prompted us to investigate the hypothesis that HIRA may regulate global chromatin architecture, thereby influencing the altered chromatin composition and spatial distribution in CML cells.

## Materials and methods

### Cell culture

K562, CML cell line was cultured using RPMI (Himedia #AL162S) with 1% Penicillin/ Streptomycin (Invitrogen #15140122) and 1% Antimycotic/Antibiotic (GIBCO# 15240-062) and 10% FBS (Gibco #10270) media at 37°C in a water jacketed incubator with 5% CO2. Cells were passaged once they achieve 70-80% confluency. Cells were stored using RPMI media +10% FBS+10% DMSO (Sigma #D2650). K562 cells are suspension cells. For distinguishing live and dead cells before seeding or storing the cells were stained with Trypan Blue stain and counted under hemocytometer.

HEK293T was cultured in standard DMEM (Himedia #AL0078) containing 10% FBS and 1% Penicillin/ Streptomycin and 1% Antimycotic/Antibiotic, as described previously (Majumder et al., 2019).

### Cloning

Cloning of H1.1 with EGFPN1 was generated using the primers mentioned in Table S2 (Fig. 2A) and used for FRAP analysis. For the DP K562 cells, H2B was amplified from H2B-mCherry and subcloned into EGFPC1 to generate H2B-EGFP for FLIM FRET analysis. The primers have been mentioned in Table S2.

### Generation of stable HIRA and SETDB1 knock down

5×10^5^ K562 cells were seeded in a 6 well plate. Next day, media changed. Added 8 µg/mL of polybrene (Sigma#H9268) to cells. Incubated at 37°C for 30 minutes. After incubation 800 µl viral particles filtered with 0.45 µ PVDF filter, were added to cells. 24 h after addition of viral particles, media was changed. Cells were selected for puromycin (Sigma#P8833) with final concentration of 1 µg/mL for three days. Knock-down was confirmed by western blot analysis. Primer sequences for shRNA mentioned in Table S3. *HIRA* shRNA was used in our earlier study (Majumder et al., 2019). *SETDB1* shRNA construct was generated in pLKO.1 vector in a similar manner as mentioned earlier (Majumder et al., 2019).

### Live cell microscopy and quantitative FRAP analysis

Live cell microscopy and quantitative FRAP analysis was done using Zeiss LSM980 Airyscan 2 microscope. Microscope is connected with an accessory incubator that maintains dark conditions as well as temperature of 37°C along with 5 % CO2 levels. For Live cell imaging, glass bottom dish (Nunc# 150680) was used. Glass bottom dish is coated with 0.01% Poly-L-Lysine (Sigma #P8920) overnight at 37 °C. Poly-L-Lysine is removed followed by wash with 1X PBS. Cells were seeded in the dish with 200 µl media and incubated at 37°C overnight. Imaging was done using 60x objective with 0.95 NA Plan apochromat. EGFP Fluorophore was excited using 488 nm solid state diode laser lines. Confocal image series were recorded with 8-bit image depth with frame size of 512 × 512-pixel size of 80 nm and with time interval of 1 seconds. For photo-bleaching, ROI of typically size of 2.5 × 1.5 µ was drawn and used for bleaching. Another ROI of similar size was drawn and used as a reference ROI either from the same cell or from a neighbouring cell of similar intensity. Background noise was normalized by using another ROI of the similar dimensions and placed at the background to calculate intensity. Only moderately intense cells were imaged for FRAP experiment. Photobleaching was done with 7 iterations 488 nm laser set at 100 % transmission. Imaging was done at 0.1% of laser intensity. For every photobleached cell, 3 pre-bleach images and post bleach frames are taken up to 120 seconds.

Analysis of the intensity calculation was done using ImageJ. To compensate the movement of the cells because of the bleaching, stackReg package was used in ImageJ. After calculating the intensity for ROI1 (Bleaching), ROI2 (Reference) and ROI3 (Background), analysis was done using following mathematical formula.

#### Formula

(Intensity of ROI before bleaching – Intensity of ROI background) / (Intensity of reference ROI after bleaching – Intensity of background ROI) Fluorescence Recovery Curve was plotted with normalized fluorescence intensity against time. Curve fitting was done with one phase association equation for linear regression. Visualization of spatial distribution of H1.1-EGFPN1 in scramble and shHIRA cells was by taking stack images at 40X apochromat at 5x zoom.

### FLIM FRET

K562 cells were co-transfected with H2B-EGFPC1 and H2B-mCherry using Lipofectamine 2000 (Invitrogen #11668019). Briefly, 2*105 K562 cells were seeded in 6 well plate. Next day, media was changed to Opti-MEM. After addition of the Opti-MEM cells were incubated at 37°C. Lipofectamine mix and DNA mix were prepared and incubated at room temperature for 5 minutes. DNA mix was added to Lipofectamine mix. Incubated for 30 minutes followed by drop-by-drop addition of the mixture to K562 cells. Post transfection 5 hours, media was changed to RPMI + 10% FBS. Cells were further selected for 14 days with 250μg/ml G418, a Neomycin analogue (Sigma #G8168) followed by FACS. K562 cells expressing both GFP and mCherry were sorted and further stored. K562 expressing H2B-EGFP and H2B-mCherry are here after referred as K562 2FP (Fluorophore positive) cells.

### Live cell imaging for FLIM-FRET

Fluorescence lifetime imaging by time-correlated single-photoncounting (TCSPC) to quantify FRET measurements – Quantifying the FLIM-FRET efficiency of each pixel allows to distinguish the euchromatin and heterochromatin spatially across nucleus. FLIM-FRET imaging was done using Leica SP8 microscope attached with TCSPC unit. Microscope was equipped with 37°C incubator that maintains 5% CO2 that allows to maintain live cells. Imaging was done using 60x oil immersion with 1.4 NA Plan-apochromat CS2 objective. Donor fluorophore EGFP was excited using 488 nm that provides 150fs pulses at an 80 MHz repetition rate. Fluorescence emission was collected at 550 nm range with mean photon count rate was adjusted to 1000 photons/second. Fluorescence lifetime was calculated for each pixel with 512 × 512 pixels or for a selected Region of interest using Leica SP8 software.

### Data Analysis for FLIM-FRET

FRET efficiency was calculated by considering the fluorescence lifetime of EGFP (Donor) in the presence and absence of the acceptor (τA). Lifetime of the donor fluorophore decreases in the presence of the acceptor. Formula used to calculate pixel wise FRET efficiency is as follows – E(FRET) = 1 - (τDA / τA),

τDA - Lifetime of the EGFP in the presence of mCherry

τD - Lifetime of the EGFP in the absence of mCherry

Mono-exponential Decay model was used to calculate the accurate fluorescence lifetime of the donor. For negating the background from the pixels with low photon count, threshold value was set. Hence low photon pixels were not considered for analysis. Χ2 was adjusted to close to 1 or lea than 1.

For calculating FRET efficiency Region of interest (ROI) was drawn with free hand tool. Bi-exponential fluorescence decay model to fit the decay curve was used to calculate fluorescence lifetime which was used to calculate FRET efficiency.

### Immunofluorescence

K562 cells were adhered to Poly-L-Lysine (Sigma-P8920) coated frosted slides by using cytospin method. Briefly 5×10^5^ cells were adhered on the slide by cytospin at 1000 RPM for 10 minutes. Cells were given a 1x PBS wash, followed by fixation with 4% PFA for 7 minutes. Cells were washed three times with 1xPBS for 5 minutes each wash. Permeabilization was done for 10 minutes using 0.2% tween20 in 1X PBS on ice. Excessive tween20 is washed with two times 1x PBS wash each for 5 minutes. 1% BSA was used for blocking. Blocking was done at room temperature for one hour. Primary antibodies were diluted in blocking buffer and incubated for overnight at 4°C (List of the antibodies Table S). Cells were washed for three times for 5 minutes each using 1X PBS. Secondary antibody labelled with fluorophore was diluted using blocking buffer and incubated at room temperature for one hour in dark. Cells were washed three times with 1X PBS to remove unbound secondary antibody. Hoechst dye was used for nuclear staining. To preserve the photobleaching, suitable mounting medium Antifade (Invitrogen #P36934) was used and coverslip was mounted on the slide. Images were acquired using Nikon A1 Rsi Confocal Laser Scanning Microscope with 60x oil immersion objective.

### Immunoprecipitation and LC-MS/MS study

K562 cells were used for pull-down of endogenous HIRA. 10 million K562 cells were pelleted and pellet was stored at −20°C. RIPA buffer was used for isolating protein. K562 cells with 2 mL of RIPA buffer (PMSF was added to RIPA buffer) were kept on ice for 30 minutes. Cells were pipetted up and down using pipette every 10 minutes. After 30 minutes incubation, cells were sonicated for 10 seconds at 25% amplitude. Followed by sonication sample was centrifuged at 12000 RPM for 5 minutes at 4°C. Supernatant was collected and used for Bradford protein estimation. Protein concentration was estimated. 10% pf the protein sample was used as an Input. For IgG control and pull-down test sample, equal amount of protein (300 µg) was used. Input was stored at −80 °C. Pre-immune serum was added to IgG sample and HIRA antibody was added to test protein sample. IgG control and protein sample were rotated at 10 RPM overnight at 4 °C on a rotator. Sepharose Beads (Sigma-P3391) were conditioned before using for IP. Sepharose beads were blocked using 2% BSA prepared in NETN buffer at 4°C for 15 minutes, followed by 3 washed with NETN buffer. IgG control and IP test samples were centrifuges at 13,000 RPM at 4 °C. Samples were added to the beads and incubated at 4 °C for 2 hours. After incubation samples were given a 5 min wash with NETN buffer four times. Finally, both IgG control and protein samples were pelleted and 2X Laemmli dye was added to it followed by heating at 85 °C for 10 minutes. After heating samples were pelleted, only supernatant was collected and stored at −20°C. Pulldown of the HIRA was confirmed by western blot analysis by probing for HIRA and ASF1A. IP sample was further processed for Mass-spectrometry. Sample was run on Poly-acryl amide gel but was not allowed to resolve. Once the sample fully runs into resolving gel, it was stained for Coomassie stain overnight, followed by destaining. Protein band from the gel was cut and put in 50 % methanol and processed for Mass spectrometry.

### Chromatin Immunoprecipitation

1×10^6^ cells per IP, were crosslinked using 1% formaldehyde for 10 minutes at room temperature followed by addition of glycine (final concentration 0.125 M). Crosslinked cells were sonicated to generate chromatin fragments. 1.5μg of either H3.3 or H3K9me3 antibodies were used to immunoprecipitate protein-DNA cross-linked fragments. Precipitated complexes were eluted and reverse crosslinked. Enrichment of chromatin fragments was measured by qRT-PCR using Sybr green fluorescence relative to a standard curve of input chromatin. IgG was used as the negative control (Majumder et al., 2019). List of primers have been enlisted in Table S4.

### ATAC-Sequencing

HIRA knockdown K562 cells were generated by shRNA guided lentiviral mediated knock-down method. Knock down was confirmed by western blot and qRT-PCR. 100000 cells of scramble and HIRA-shRNA were pelleted and used for ATATC-sequencing. ATAC-sequencing was done according to standard protocol followed by Active Motif.

### EdU incorporation assay

To trace the replication status upon downregulating HIRA in K562 cells, we employed the click chemistry mechanism by pulsing the cells with 5µM alkyne conjugated nucleoside analogue, EdU (5-ethynyl-2’-deoxyuridne) (Thermo Fisher Scientific, #A10044) for 4 hours after incubating cells with serum-deprived media (RPMI +10% FBS) for 16 hours, followed by providing the complete medium (RPMI+ 10%FBS) for 2 hours. This step was followed by coupled fixation and permeabilization with fix buffer A (300mM sucrose, 0.5% Triton-X-100 and 2% formalin), followed by blocking with 10% FBS containing PBS. We performed a copper-catalyzed azide coupling reaction that has 10µM Alexa fluor 488 azide (Invitrogen, #A10266) in the dark for 1 hour, at room temperature. Cells were incubated with propidium iodide staining solution after washing with 1X PBST and subjected to FACS analysis.

### Protein isolation and Immunoblotting

Protein was isolated from freshly pelleted cells using Radioimmunoprecipitation (RIPA) buffer containing using sonication. RIPA buffer contains (10mM Tris-HCl, pH7.5, 150Mm NaCl, 5mM EDTA, 1% Triton X-100, 1% Nonidet P-40, 1% Sodium deoxycholate, 0.1%SDS, 1mM Sodium orthovanadate, and 1mM PMSF) (Majumder et al., 2015). Isolated protein concentration was estimated by Bradford Reagent (Bio-Rad #500-0006). By loading equal quantity of proteins on different percentage of gels as per requirements 8%, 10%, 12%, protein was resolved on PAGE followed by transfer it on PVDF membrane (Merck IPVA00010) followed by 1 hours blocking with 5% skimmed milk. Antibodies used have been enlisted in Table S5. Visualization of the protein was by enhanced chemiluminescence.

### Quantitative RT-PCR

RNA was isolated using RNAeasy kit (Qiagen #74106) following the manufacturers guidelines. cDNA synthesis was aided by random hexamer along with Reverse transcriptase enzyme (ABI Biosystems #4368814). qRT-PCR was performed with SYBR-Green (ABI Biosystems #4309155) and analyzed by using relative standard curve method. Primers used are listed in Table S6.

## Results

### HIRA associate with heterochromatin in CML cells

HIRA regulate chromatin accessibility during hematopoiesis, but how it could modulate the chromatin during leukemia remains unexplored. We investigated upon the HIRA interactome in CML cells. K562, the CML cell line, served as the model for the entire study. Whole cell protein lysate from K562 cells were subjected to immunoprecipitation with HIRA followed by LC/MS-MS analysis. The resulting HIRA-pulled down lysate was confirmed with presence of HIRA and histone variant H3.3, the bonafide HIRA interacting partner (Fig. 1A). Using NetworkAnalyst bioinformatics tool (Zhou et al., 2019), the generic protein-protein interaction network for HIRA was generated (Fig. 1B). The top-most enriched KEGG pathway for HIRA was carcinogenesis, followed by cell cycle, apoptosis, cellular senescence among others (Fig. 1B, Table S1). A total of 49 hits were obtained for HIRA. DAVID bioinformatics tool analysis demonstrated enrichment of nucleosome and heterochromatin components among top 10 biological processes (Fig. 1C, red box). Interestingly, the study conducted by Radich et al. (2006) indicated association of HIRA during the progression of CML from chronic to blast phase of the disease (Radich et al., 2006, supplementary table 8 has been referred here). To demonstrate whether an alteration in HIRA expression could influence the expression of genes implicated during the progression of leukemia, we knocked down HIRA in CML K562 cells (*HIRA-*shRNA) (Fig. 1D). CML is distinguished by the Philadelphia (Ph) chromosome, which arises from balanced reciprocal translocation (t(9;22)(q34;q11) resulting in the expression of the BCR-ABL1 oncogene, which encodes the chimeric BCR-ABL1 protein with constitutive kinase activity. The expression of BCR-ABL1 is absolutely essential for the maintenance and also for the progression of the disease (Quintás-Cardama and Cortes, 2009). Eventually, BCR-ABL1 modulates the expression of transcription factors associated with differentiation, resulting in its arrest (Carlesso et al., 1996). Upon loss in HIRA, BCR-ABL1 level was downregulated both at the protein and RNA level (Fig. 1D, 1E). Induced level of the components of Wnt signaling pathway has been reported to be significantly associated with the progression of CML along with increase in BCR-ABL expression (Radich et al., 2006). Interestingly, upon knock-down of HIRA, we observed a downregulation of E-cadherin (CDH1), a component of Wnt signaling pathway and BCR-ABL expression at the mRNA level. In addition to that CML disease progression also associate with loss in myeloid differentiation, apoptosis and DNA damage repair (Radich et al., 2005). However, upon downregulation of HIRA, we observed a significant increase in the level of myeloid differentiation marker CCAAT Enhancer Binding Protein Alpha (*CEBPA*) (Quintás-Cardama and Cortes, 2009), apoptosis marker B-cell lymphoma-2 (*BCL2*) and DNA damage repair marker 5’-3’ Exoribonuclease 2 (*XRN2*) (Fig. 1E). This outcome highlighted the importance of investigating the role of HIRA in regulation of CML and additionally also hinted that HIRA might have a role in the progression of the disease from chronic to blast phase.

**Figure 1.**
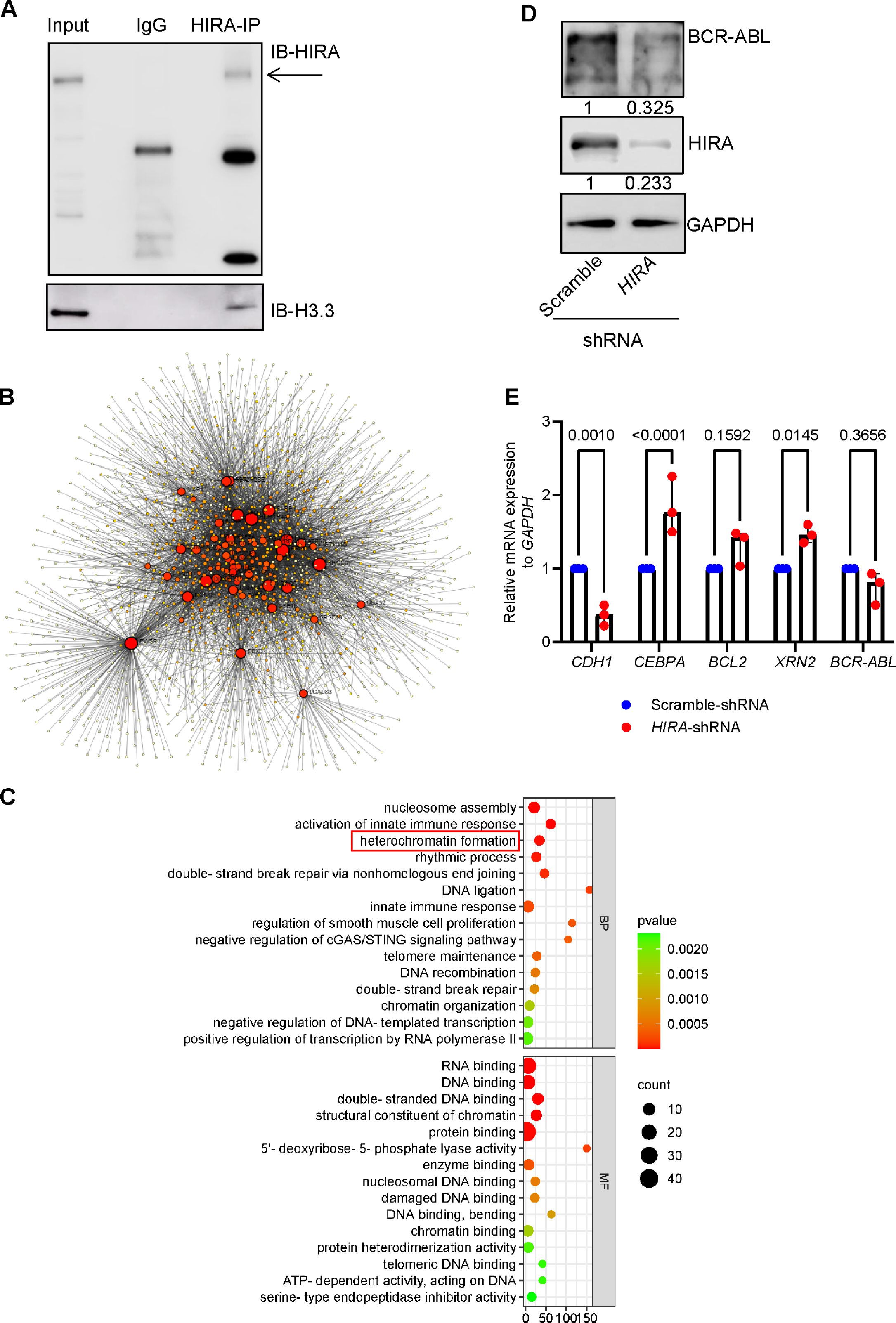
Human HIRA interactome in CML cells. A. Western blot analysis for the confirmation of immunoprecipation of K562 cell lysate with HIRA. Interaction of H3.3 with HIRA. Total protein was isolated from K562 cells. HIRA interacting proteins were pulled down using HIRA antibody by immunoprecipitation and the complex extracted from Protein-A sepharose beads was run on a SDS-PAGE and the gel piece was subjected to LC/MS-MS analysis for the determination of interacting partners of HIRA. B. The protein-protein interaction network for HIRA was deduced by NetworkAnalyst (refer to Table S1 for the KEGG pathways). C. Bubble graph demonstrate the Gene ontology (GO) analysis by DAVID bioinformatics tool (https://david.ncifcrf.gov) for the top 15 biological processes (BP) and molecular function (MF) for HIRA. D. Western blot analysis for the expression of HIRA and fusion BCR-ABL in scramble and *HIRA*-shRNA expressing K562 cells. Lentiviral particles harbouring *HIRA*-shRNA or scramble shRNA were transduced in K562 cells, followed by selection in presence of puromycin. Cell lysate was isolated and subjected to SDS-PAGE. E. Bar graph represent qRT-PCR analysis for the expression of genes associated with progression of CML in the same set of cells analyzed in D. Unpaired t-test was performed for the statistical analysis, p<0.05 is significant.

But, how HIRA drives this change in the behavior of CML cells? Being a histone chaperone, we investigated upon the role of HIRA in regulation of the chromatin status of these cells.

### HIRA regulate chromatin compaction

Chromatin is highly dynamic and it gets compacted and de-compacted as per the requirements of the cell physiology. To understand the role of HIRA in modulating chromatin architecture in CML cells, we compared the global euchromatin and heterochromatin content of the wild type and HIRA depleted K562 cells using Fluorescence Recovery After Photobleaching (FRAP). Histone H1.1 being a linker histone is mobile in loosely compacted chromatin (i.e. euchromatin) while its mobility gets restricted in a densely packed or compacted chromatin (i.e. heterochromatin) (Kirmes et al. 2015; Lever et al. 2000; Long et al. 2021). Hence, we tagged GFP at N-terminus of H1.1 for FRAP analysis. The GFP-positive cells were sorted by FACS (Supplementary figure S1A-S1C). Stable K562 cells expressing H1.1-EGFPN1 were generated followed by downregulation of HIRA by lentiviral mediated shRNA expression (Fig. 2A, 2B, 2C). Interestingly live cell imaging of the scramble K562 cells showed uniformly distributed H1.1-EGFPN1 within the nucleus, on the contrary HIRA knockdown K562 cells showed H1.1-EGFPN1 significantly localized at the nuclear periphery (Fig. 2D, 2E Additional video 1 and 2). FRAP analysis demonstrated that the ROI bleached either at the nuclear periphery or in the middle of the nucleus showed up to 80% recovery in scramble-shRNA cells indicating the euchromatin state while cells expressing *HIRA*-shRNA could recover only up to 40% of the fluorescence (Fig. 2F, 2G). Striking difference in the recovery rate of H1.1-EGFPN1 indicates that upon down-regulation of HIRA there is increase in the heterochromatin content accompanied by its re-distribution towards the nuclear periphery (Fig. 2C-2G). Significant increase in the half time for fluorescence recovery after photobleaching was observed in cells transfected with *HIRA*-shRNA compared to scramble-shRNA (Fig. 2H). On the contrary, a significant decrease in mobile fraction was observed in the *HIRA*-shRNA expressing cells in comparison to the scramble-shRNA expressing cells (Fig. 2I). Interestingly, upon rescue of the *HIRA*-shRNA cells with ectopic expression of *HIRA*-Flag, we observed a significant increase in the recovery of the fluorescence, increase in the mobile fraction and reduction in the half time of recovery (Supplementary figure S1D, S1E, S1F). This substantiated the fact that loss in expression of HIRA indeed contributes to the compaction of the chromatin in leukemia cells.

**Figure 2.**
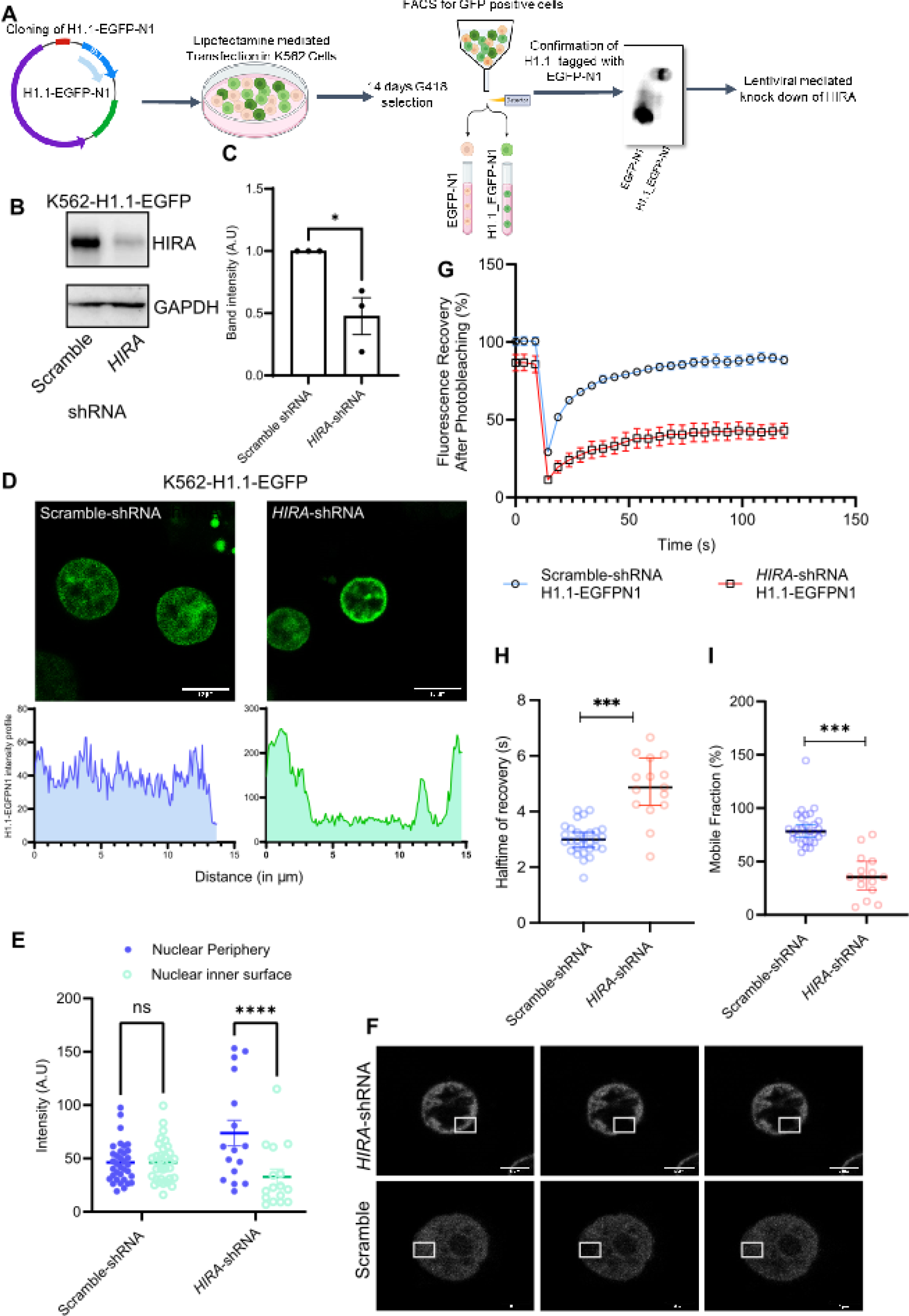
HIRA regulate chromatin compaction in CML cells. A. Schematic representation of FRAP assay. B. Western blot analysis for the expression of HIRA in H1.1-EGFP positive scramble and HIRA-shRNA expressing K562 cells. C. Band intensity for the knockdown of HIRA in H1.1-EGFP K562 cells. D. Live cell imaging for the expression of GFP in the same set of cells analyzed in B. Lower panel demonstrate the intensity curve of GFP across the nucleus. E. Scatter plots represent the intensity of GFP at the nuclear periphery and within the inner surface of the nucleus in the same set of cells analyzed in B. A two-way ANOVA statistical analysis was done followed by Šídák’s multiple comparisons tests, ****p<0.0001, ns=not significant. F. FRAP analysis in K562 cells. The micrographs represent ROI (square box) for pre-bleach, bleach and post-bleach area. G. Fluorescence recovery curve of the same set of cells. H, I. Scatter plots represent the mobile fractions and half time for recovery in scramble and *HIRA*-shRNA expressing cells. Two-tailed unpaired t-test was performed for the statistical analysis, ***p<0.001.

Next, we were intrigued to determine whether the reduction in HIRA expression levels was causing a spatial redistribution of chromatin, as live cell imaging of the cells revealed a shift in chromatin positioning.

### HIRA regulate spatial distribution of chromatin

FRAP analysis gave an insight to the role of HIRA in modulating chromatin architecture. In order to quantify the chromatin compaction induced by depletion of HIRA and visualize the spatial distribution across the nucleus, we employed Fluorescence lifetime imaging microscopy (FLIM)- Forster Resonance Energy Transfer (FRET). H2B tagged with donor (EGFP) and acceptor (mCherry) fluorophore separately were stably transfected in K562 cells followed by the sorting of double positive (DP) cells (Supplementary figure S1G). These cells were subjected to the knockdown of HIRA using lentiviral mediated shRNA expression (Fig. 3A-3D). FLIM-FRET allows to distinguish the euchromatin and heterochromatin based on the lifetime of the donor fluorophore H2B-EGFP. H2B-EGFP and H2B-mcherry has been previously used as a FRET pair for FLIM-FRET analysis (Llères et al. 2009, 2017). When chromatin is compacted H2B-EGFP (Donor fluorophore) and H2B-mCherry (acceptor fluorophore) being in close proximity, lead to FRET, consequently reducing the lifetime of the fluorophore. Due to the difference in compaction, heterochromatin will show increased FRET efficiency while euchromatin will show less FRET efficiency. Pixel wise counting of the donor lifetime allows calculating FRET efficiency, which represents the state of chromatin. Upon down-regulation of HIRA in DP cells, a re-organization of chromatin was observed. FRET efficiency map of HIRA down-regulated cells showed similar observation from FRAP with spatial redistribution of the chromatin at the nuclear periphery and increase in compaction or enhanced presence of heterochromatin (Fig. 3E). Loss in HIRA led to significant decrease in lifetime of the EGFP in comparison to the control cells (Fig. 3F). Upon rescue of the HIRA level by ectopic expression of HIRA-Flag in the DP cells, the FRET-efficiency was reduced with redistribution of the chromatin throughout the nucleus (Supplementary figure S1H). FLIM-FRET analysis demonstrated that HIRA downregulation leads to increase in the heterochromatin content associated with alteration in the spatial distribution of the chromatin. Does the alteration in chromatin architecture could influence overall accessibility of the chromatin?

**Figure 3.**
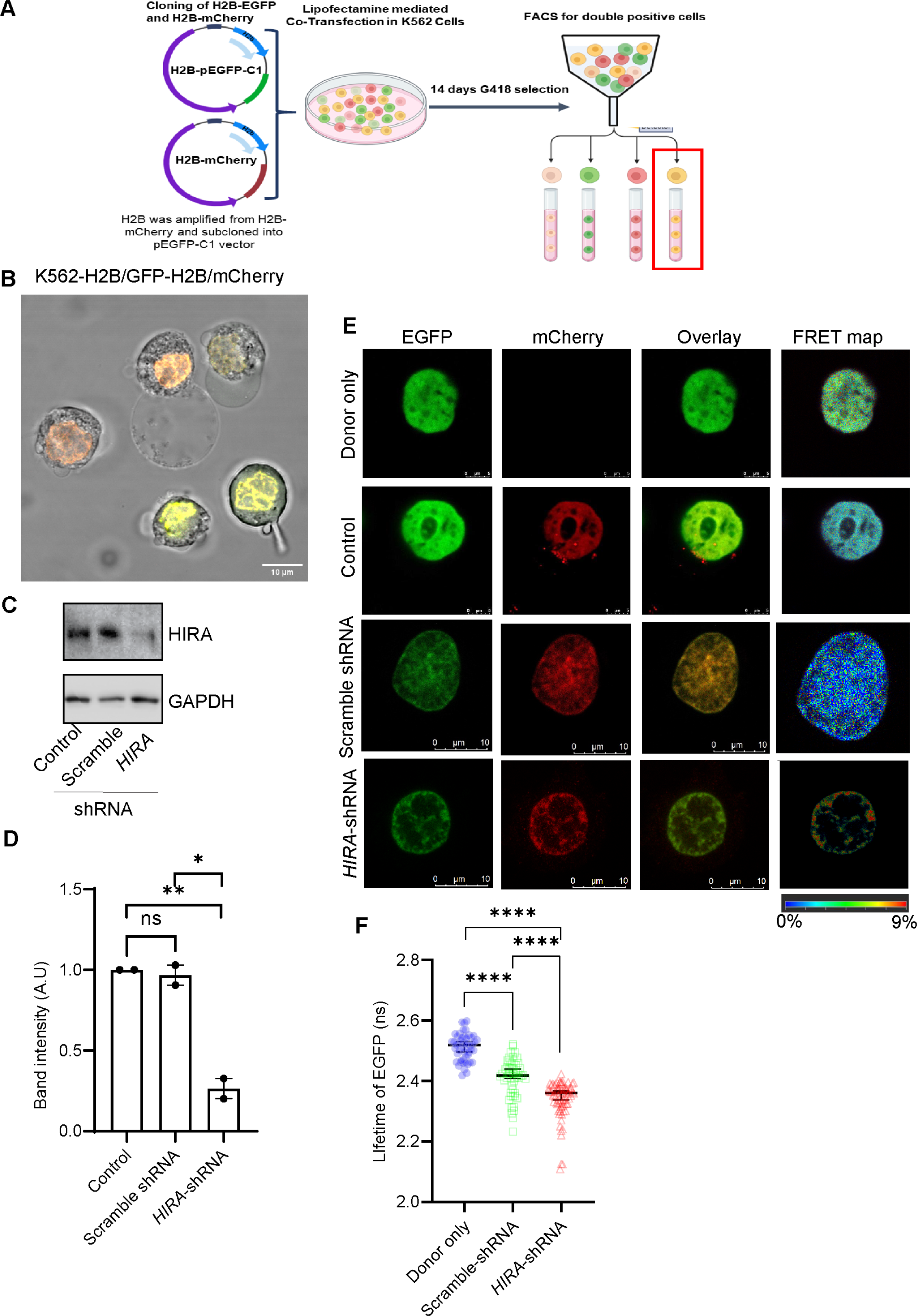
HIRA regulate spatial distribution of the chromatin in CML cells. A. Schematic representation of FLIM-FRET assay. B. Live cell imaging for the co-expression of GFP and mCherry in K562 cells. C. Western blot analysis for the expression of HIRA in double positive control, scramble and *HIRA*-shRNA expressing K562 cells. D. Bar graph represents the band intensity for the knockdown of HIRA in double positive K562 cells. Unpaired t-test was performed for the statistical analysis, *p<0.05, **<p<0.01, n=2 biological replicates. E. Live cell images for the donor-acceptor and FRET efficiency map. F. Scatter plot demonstrate the significant reduction in life-time of GFP in the same set of cells analyzed in C. Two-tailed unpaired t-test was performed for the statistical analysis, ***p<0.001.

### HIRA regulate global chromatin accessibility in leukemia cells

To understand the role of HIRA on the overall change in the accessibility of chromatin, we performed ATAC-sequencing of the chromatin derived from K562 cells expressing scramble or *HIRA*-shRNA. Upon downregulation of HIRA, a loss in accessibility was observed within the promoter region (−5kb to +5kb) (Fig. 4A) and the gene bodies (Fig. 4B). Next, we were intrigued to investigate the chromatin accessibility within the genomic loci linked to CML or its progression from the chronic to the blast phase. Radich et al. (2006) formulated the top ten genes that are upregulated or downregulated in CML, irrespective of the CD34 status (Radich et al., 2006, supplementary Table 8: Top 10 genes in progression set has been referred here). Upregulated genes (*GCL2, PRAME*) in CML progression demonstrated a reduced accessibility upon loss in HIRA level (Fig. 4C, 4D, Supplementary figure S2A-S2D). Reduced chromatin accessibility indicates a downregulation in the expression of these genes. On the other hand, downregulated genes (EREG, CXCL8) during disease progression, demonstrated an increase in accessibility upon loss in HIRA level (Fig. 4E, 4F, Supplementary figure S2E-S2H) and this indicate an upregulation in the expression of the genes. When we investigated into the status of the progression genes (Radich et al., 2006, Table 1: Functional annotation of “progression” genes, have been referred here) a similar inverse trend was observed upon loss in expression of HIRA (Fig. 4G-4L). Ribosome (RPL13A) and Wnt signaling (CDH1) related genes showed reduced accessibility of the chromatin, whereas myeloid differentiation (CEBPE), nucleosome (BAZ1A), apoptosis (BCL2) and DNA damage response (XRN) associated genes showed enhanced accessibility within the locus (Fig. 4G-4L). This further validate the alteration in expression of genes at the mRNA level mentioned in Fig. 1E. Although, no effect on genes associated with sugar metabolism was observed upon downregulation of HIRA. This finding indicates the role of HIRA and implies its significance in the disease progression from chronic to blast phase.

**Figure 4.**
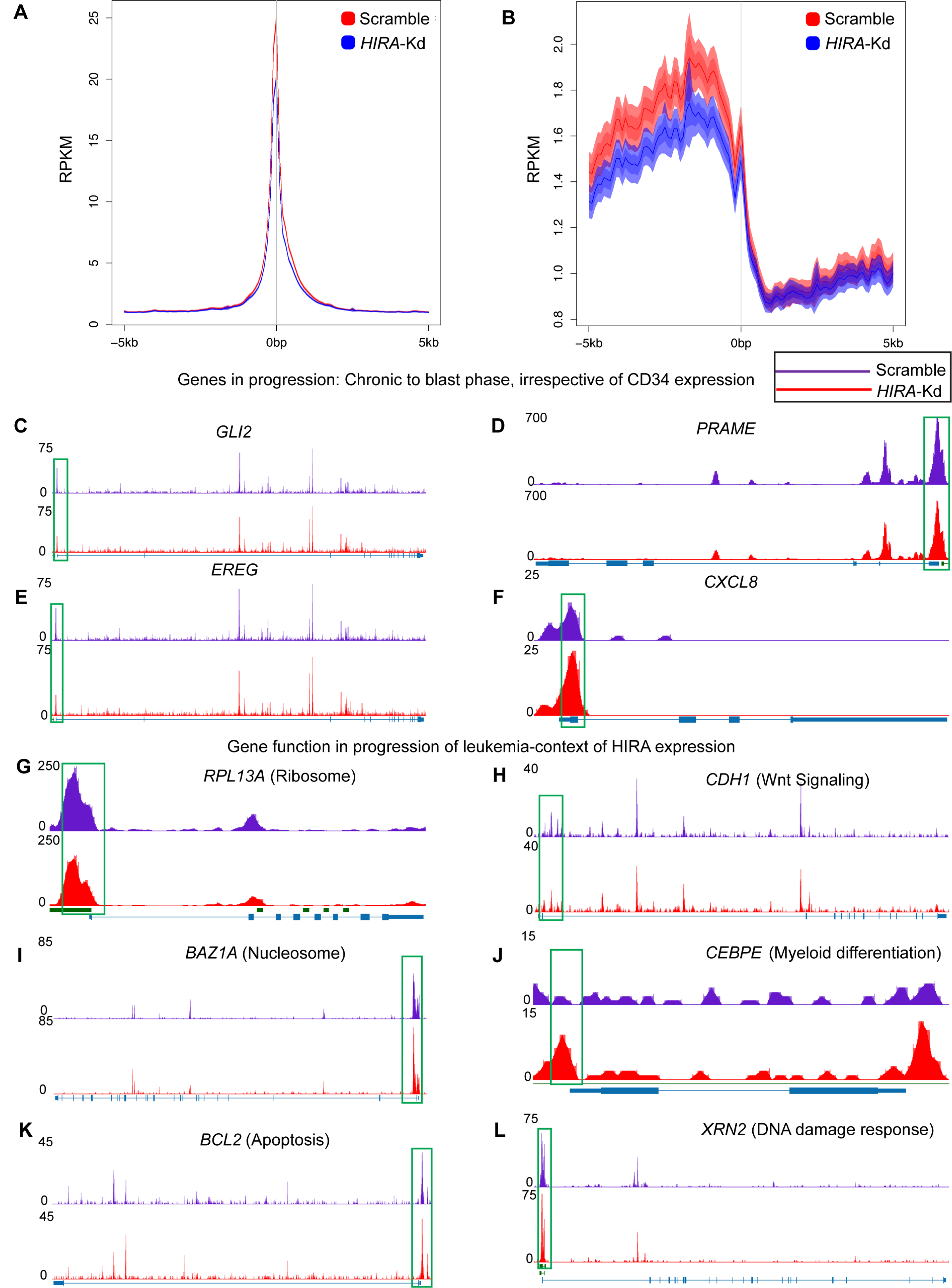
HIRA regulate global chromatin accessibility in leukemia cells. A, B. Average profiles of chromatin accessibility in scramble-shRNA and *HIRA-*shRNA (mentioned as HIRA-kd) expressing K562 cells at pooled ATAC-seq peaks. The y axis represents the RPKM values, and the x axis represents distance from the center (kb). C-E. Representative UCSC tracks showing the ATAC-seq profiles for genes (*GLI2, PRAME*) with decreased chromatin accessibility in the promoter regions (C, D) and genes (*EREG, CXCL8*) with increased accessibility (E, F) in the progression of CML (Radich et al., 2006). G-L. UCSC tracks showing the ATAC-seq profiles for genes representing different GO with decreased chromatin accessibility in the promoter regions of ribosome (*RP13A*), Wnt signaling (*CDH1*) (G, H) and genes with increased accessibility in the promoter regions of nucleosome (*BAZF1*) (I), myeloid differentiation (*CEBPE*) (J), apoptosis (*BCL2*) (K) and DNA damage response (*XRN2*) (L) in the progression of CML from chronic to blast stage. Green boxes indicate the region within the locus (promoter region) showing alteration in peak heights. The ATAC-sequencing was performed at Active Motif, USA.

But how loss of HIRA could result in chromatin compaction, altered spatial distribution and reduced chromatin accessibility in the CML cells?

### Downregulation of HIRA induce global increase in H3K9me3 level

H3.3 histone variant is a bonafide interacting partner of HIRA and in K562 cells, the same association was maintained (Fig. 1A). But, here we observed an increase in heterochromatin upon downregulation of HIRA. Hence, we investigated into the global change in histone modification patterns that could result in increased heterochromatin formation. We determined the level of histone H3K4me3, H3K4me2, H3K4me1, H3K4ac, H3K9ac, H3K27me3, H3K9me3 levels in response to the loss in expression of HIRA (Fig. 5A). Interestingly, among all of them, a significant increase was observed in the level of H3K9me3 only (Fig. 5B). Immunofluorescence analysis indicated the presence of H3K9me3 at the nuclear periphery in HIRA-knockdown cells (Fig. 5C). It can be recalled here that a similar expression of H1.1-EGFP and spatial distribution of chromatin was observed under similar condition (Fig. 2, Fig. 3). Thus, it’s the increase in the level of H3K9me3 that is contributing to the enhanced compaction of the chromatin.

**Figure 5.**
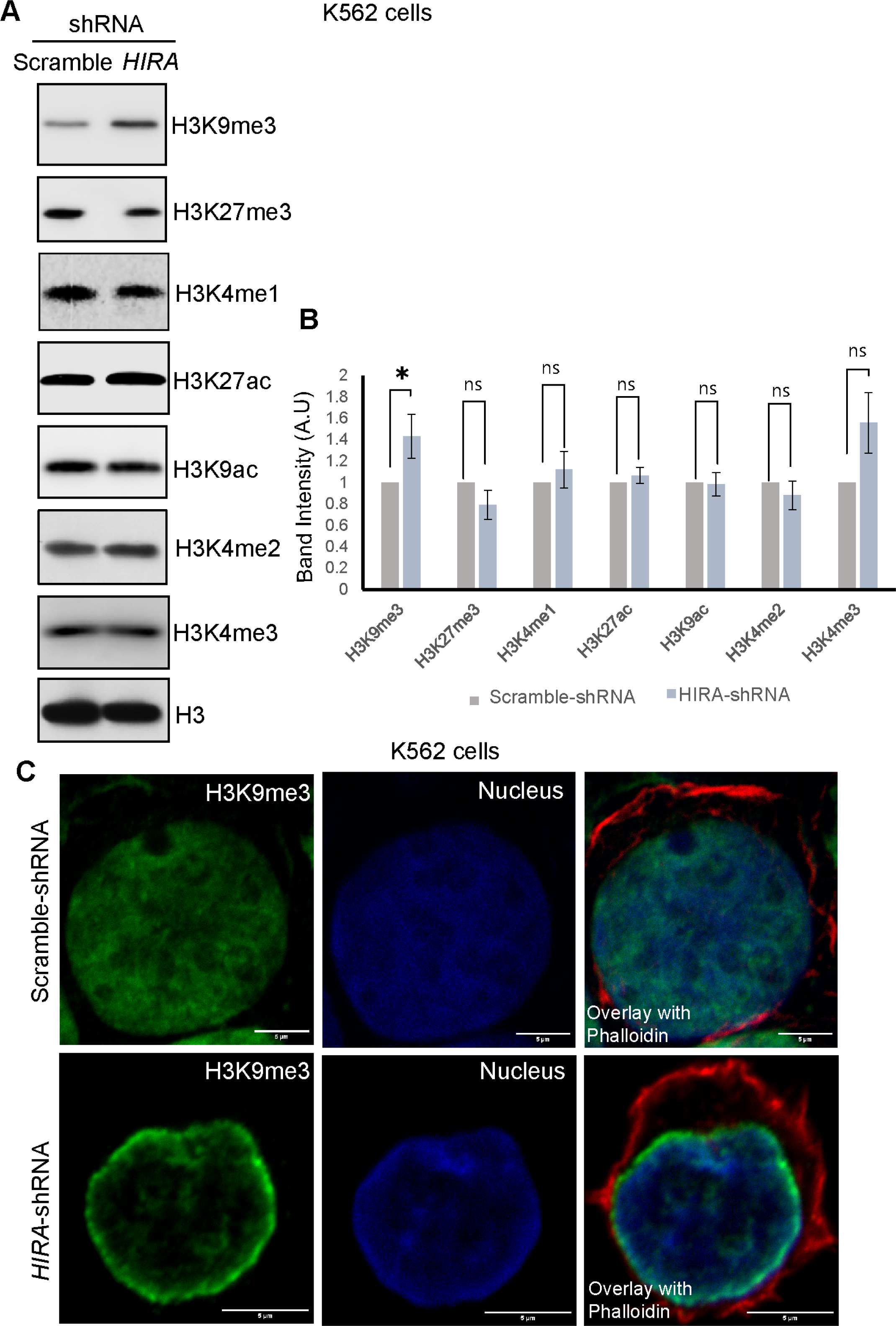
Loss in HIRA induce global increase in H3K9me3 level. A. Western blot analysis for the expression of different histone modification marks in scramble vs *HIRA*-shRNA expressing K562 cells. B. Bar graph representing the band intensities of histone modification marks. A two-way ANOVA statistical analysis was done followed by Šídák’s multiple comparisons tests, *p<0.05, ns=not significant. C. Immunofluorescence analysis for the expression of H3K9me3 in scramble vs *HIRA*-shRNA expressing K562 cells.

But, how does the level of H3K9me3 is altered upon loss in HIRA?

### HIRA-SETDB1-H3K9me3 axis regulate chromatin architecture in CML cells

The addition or removal of functional group from the lysine residue of the histone tails are accomplished by a set of histone modifying enzymes, the demethylase or the methyltransferases. Hence, to investigate upon the fact that which mechanism drive the increase in H3K9me3 level, we determined the level of these modifying enzymes. Interestingly, among the demethylases, we failed to observe any significant change upon loss of HIRA expression (Fig. 6A), however, a significant increase in SETDB1 (SET Domain Bifurcated Histone Lysine Methyltransferase 1) level was observed both at the RNA and protein level (Fig. 6A, 6B). But, does this increase in SETDB1 expression really contribute to the increase in H3K9me3 level? To answer that, we knocked down SETDB1 in *HIRA*-knockdown K562 cells (Fig. 6B) and observed the level of H3K9me3. The increase in H3K9me3 level was abrogated with loss in SETDB1 level (Fig. 6B). Thus, SETDB1 mediate the increase in H3K9me3 level in the context of loss of HIRA in K562 cells. But, how does altered HIRA level enhance SETDB1 expression? Incorporation of H3.3 mediated by HIRA complex and/or DAXX/ATRX within the chromatin, dictate contradictory functions in the context of development and differentiation (Choi et al., 2024). We determined the incorporation or loss of H3.3 in the scramble and *HIRA*-shRNA expressing K562 cells within the *SETDB1* promoter. Interestingly, we observed a significant enrichment in the incorporation of H3.3 within the *SETDB1* promoter upon downregulation of HIRA (Fig. 6C). This finding underlies the enhanced expression of SETDB1 in K562 cells expressing *HIRA*-shRNA. The K562 cells generated for FRAP analysis (Fig. 2) were further manipulated for the downregulation of SETDB1, leading to the generation of HIRA/SETDB1-double knockdown cells. Live cell imaging of HIRA-downregulated cells demonstrated similar chromatin architecture with enhanced presence in nuclear periphery (Fig. 6D) as mentioned in figure 2 whereas upon downregulation of SETDB1, these cells demonstrated the chromatin distribution throughout the nucleus, like the one present in cells expressing only scramble-shRNA (Fig. 6E). FRAP analysis indicated increase in recovery of fluorescence upon loss in SETDB1 expression in HIRA-downregulated cells (Fig. 6F). Similarly, an increase in mobile fraction and decrease in halftime in recovery was observed in the HIRA/SETDB1 knocked down cells. Thus, we provide evidence that the SETDB1 expression was responsible for the enhanced H3K9me3 level upon loss in HIRA that aid in the regulation of chromatin architecture of CML cells.

But, how does this chromatin architecture regulate cellular function of CML cells?

**Figure 6.**
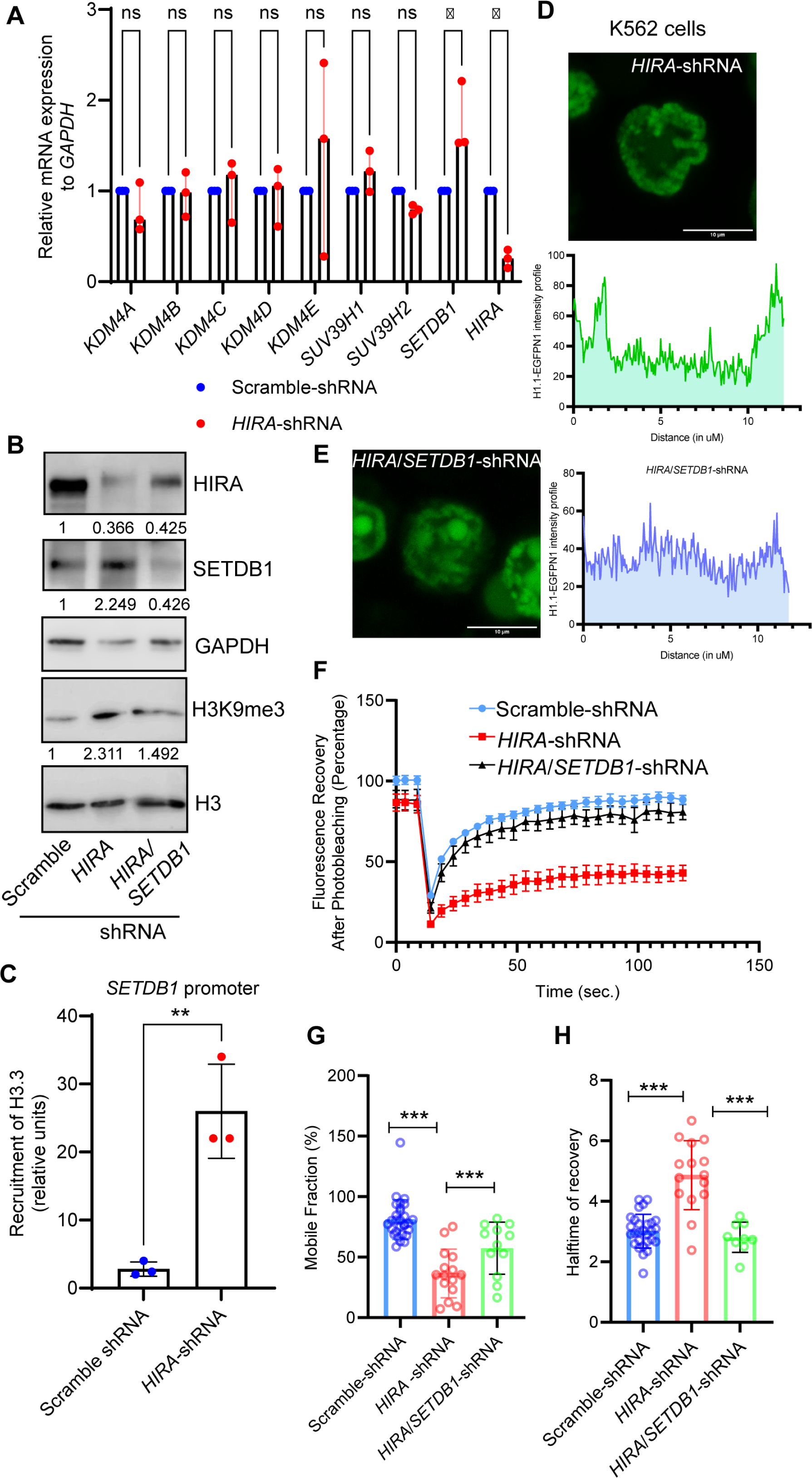
HIRA-SETDB1-H3K9me3 axis regulates chromatin architecture in CML cells. A. Quantitative RT-PCR analysis for the expression of different histone methyltransferases and demethylases in scramble vs *HIRA*-shRNA expressing K562 cells. A two-way ANOVA statistical analysis was done followed by Šídák’s multiple comparisons tests, *p<0.05, ns=not significant, n=3 biological replicates. B. Western blot analysis for the expression of SETDB1, HIRA, H3K9me3 level in scramble vs *HIRA*-shRNA expressing K562 cells. C. Bar graph represents the quantitative ChIP analysis for the recruitment of histone variant H3.3 within *SETDB1* promoter in scramble vs *HIRA*-shRNA expressing K562 cells. Two-tailed unpaired t-test was performed for the statistical analysis, **p<0.01, n=3. D, E. Live cell imaging for the expression of GFP in the HIRA-shRNA and HIRA/SETDB1-shRNA expressing K562 cells. Intensity curve of GFP across the nucleus has been shown. F. Fluorescence recovery curve of the same set of cells analyzed in D, E. G, H. Scatter plots represent the mobile fractions and half time for recovery in scramble, *HIRA*-shRNA and HIRA/SETDB1-shRNA expressing cells. Two-tailed unpaired t-test was performed for the statistical analysis, ***p<0.001.

### Altered chromatin architecture influence proliferation vs differentiation of CML cells

Our earlier study demonstrated that loss in HIRA could dictate proliferation vs differentiation (Majumder et al., 2019). Here also, we performed 5-Ethynyl-2′-deoxyuridine (EdU) incorporation assay to further confirm the role of HIRA in proliferation of CML cells. Upon downregulation of HIRA, we observed a significant reduction in EdU incorporation in K562 cells (Fig. 7A, 7B). This further consolidates our earlier finding that loss in HIRA inhibit proliferation. But does loss in HIRA influence the expression of proteins associated with cell proliferation by inducing an alteration in their respective chromatin architecture? How does HIRA mediate it? ATAC-seq analysis revealed a reduction in chromatin accessibility within proliferation markers including and *PCNA* and *MiK67* loci upon knockdown of HIRA (Fig. 7C, 7D). Quantitative ChIP analysis demonstrated an increased abundance of heterochromatin mark H3K9me3 within the respective promoters in HIRA downregulated cells in comparison to control cells (Fig. 7E, 7F). When we investigated into the genes associated with erythroid or megakaryocytic differentiation, we observed a differential chromatin accessibility status and a dynamic pattern for H3K9me3 incorporation within these promoter regions (Fig. 7G-7L). The chromatin accessibility was reduced within the erythroid specific cell surface marker (*GYPA and TFRC*) (Fig. 7G, 7H), while an increased accessibility was observed within megakaryocyte specific cell surface marker (*GP6 and GPIIB*) (Fig. 7I, 7J). We observed an increase in the incorporation of H3K9me3 level within erythroid-specific *GYPA* promoter while an opposite trend was observed within the megakaryocyte-specific *GP3A* promoter (Fig. 7K, 7L). Interestingly, this induction in megakaryocyte differentiation was abrogated in HIRA/SETDB1 downregulated K562 cells. Increased MKL1 and GATA2 expression in HIRA-downregulated cells demonstrated a reduced expression upon loss in SETDB1 expression in the same cells (Fig. 7M). This further establishes the role of SETDB1 in the HIRA mediated regulation of chromatin in CML cells. Thus, HIRA-SETDB1-H3K9me3 axis might modulate the proliferation vs differentiation of CML cells in a context specific manner.

**Figure 7.**
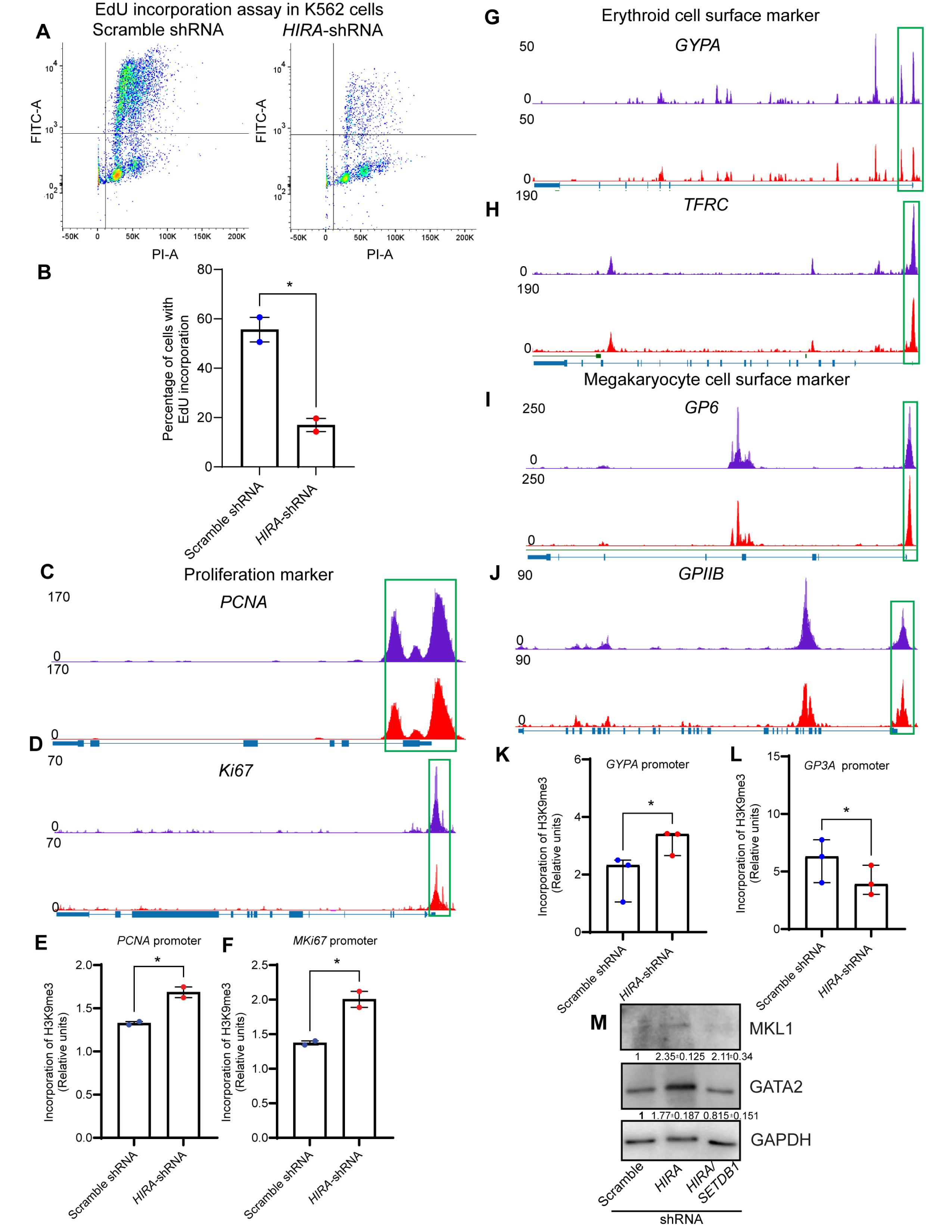
HIRA mediate altered chromatin architecture dictating proliferation vs differentiation. A, B. FACS analysis for the quantification of EdU positive population in K562 cells expressing scramble and HIRA-shRNA. Bar graph represent the incorporation of EdU in the cells. Two-tailed unpaired t-test was performed for the statistical analysis, *p<0.05, n=2. C, D. Representative UCSC tracks showing the ATAC-seq profiles for cell proliferation associated genes (*PCNA, Mki67*) with decreased chromatin accessibility in the promoter regions upon downregulation of HIRA. E, F. Bar graph represent the quantitative ChIP analysis for the incorporation of histone H3K9me3 mark within the promoter region of Ki67 and PCNA. Two-tailed unpaired t-test was performed for the statistical analysis, *p<0.05, n=2. G, H. Representative UCSC tracks showing the ATAC-seq profiles for genes (*GYPA*, *TFRC*) with decreased chromatin accessibility in the promoter regions associated with erythroid differentiation. I, J. Representative UCSC tracks showing the ATAC-seq profiles for genes (*GP6, GPIIB*) with increased chromatin accessibility in the promoter regions associated with megakaryocyte differentiation. Green boxes indicate the region within the locus (promoter region) showing altered presence of peaks. ATAC-sequencing was performed at Active Motif, USA. K, L. Bar graph represent the quantitative ChIP analysis for the incorporation of histone H3K9me3 mark within the promoter region of *GYPA* and *ITGB3*. Two-tailed unpaired t-test was performed for the statistical analysis, n=3. M. Western blot analysis for the expression of megakaryocyte specific markers (MKL1, GATA2) in cells expressing scramble-shRNA, *HIRA*-shRNA and *HIRA/SETDB1*-shRNA.

Thus, here in this study we provide experimental evidences that demonstrated that histone chaperone HIRA could regulate the expression of histone methyl transferase SETDB1, thereby regulating the chromatin architecture in terms of compaction, spatial distribution and accessibility by incorporation or loss of heterochromatin mark H3K9me3, which in turn could regulate the balance between proliferation vs differentiation of CML cells (Fig. 8).

**Figure 8.**
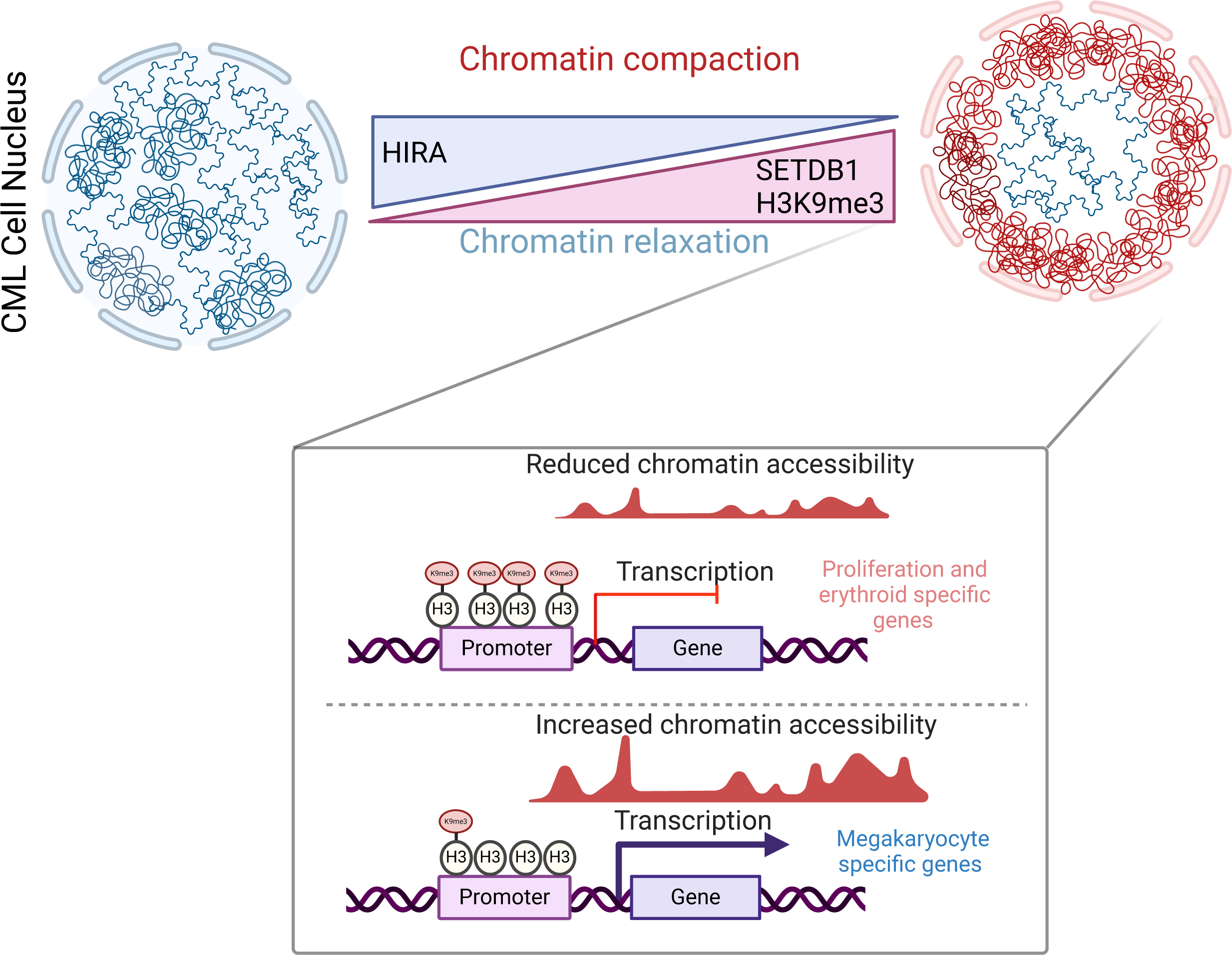
Model depicting the regulation of chromatin architecture in CML cells by HIRA. The model was generated using Biorender.

## Discussions

Histone chaperone HIRA has been implicated in multiple developmental processes, senescence and tumor growth with its knockout phenotype resulting in early embryonic lethality (Robert et al., 2002). Patients with DiGeorge syndromes have a deletion region including a stretch of *HIRA* locus (Chen et al., 2020). These patients suffer from immunodeficiency along with thrombocytopenia. But for a long time, the understanding of HIRA in the context of hematopoiesis was not explored.

Lately, reports from our lab and others have demonstrated the role of HIRA in hematopoiesis (Majumder et al., 2019; Chen et al. 2020; Cabal-Hierro et al., 2020). When hematopoietic stem cells acquire genetic or epigenetic aberrations, the normal hematopoiesis gives rise to the cancer of the blood, which is leukemia. Restricted differentiation and uncontrolled proliferation mark leukemia condition. Our recent study showed the enhanced expression of HIRA in CML patients vs normal or AML patients and CML cell line K562 showed maximum enrichment of HIRA as per CCLE analysis (Majumder et al., 2019). But, till date very limited number of studies was conducted to understand the role of HIRA in leukemia. Especially when, in the 2006 study by Radich et al., HIRA was among the embryonic development cluster that associated with the progression of leukemia from chronic to blast phase (Radich et al., Supplementary Table 9 is referred). Chronic myeloid leukemia, a hematopoietic stem cell disease, progresses from the chronic phase through the accelerated phase to the blast phase. Distinct gene expression pattern guide this 2-step progression process.

Our analysis of the HIRA-downregulation in CML cells showed loss in expression of CDH1 along with reduced accessibility of the chromatin, one of the components of Wnt-signaling pathway which is potentially significant in the progression of CML. Basically, out of the 7 major functionally annotated groups (Radich et al., 2006), except for sugar metabolism, an altered chromatin accessibility was observed upon downregulation of HIRA within representative genes of all 6 groups. These observations provide significant importance to understand the role of HIRA in leukemia and its progression.

HIRA interactome demonstrated its interaction with proteins associated with heterochromatin. Earlier reports provided evidence on role of HIRA in the formation of senescence-associated heterochromatin foci (Zhang et al., 2005). But how it is associated with leukemia has not been studied. HIRA selectively incorporates histone variant H3.3 in a replication independent manner into the nucleosomes within defined chromatin domains. But the incorporation of H3.3 has contradictory roles at different loci. In fact, the H3.3-HIRA complex is responsible for the formation of heterochomatin in senescent cells. We observed that upon loss in HIRA, chromatin compaction, spatial distribution and accessibility was altered in the CML cells. But this must have been quite evident, as deprivation of HIRA led to reduced H3.3 incorporation leading to repressed chromatin. But the spatial redistribution of the chromatin at the nuclear periphery indicated an additional factor and that is H3K9me3 deposition, which is a bonafide heterochromatin mark.

ATAC-sequencing of the HIRA-deprived chromatin in CML cells, indicated both the reduced as well as enhanced chromatin accessibility. As mentioned earlier, the incorporation of H3.3 or H3K9me3 is context specific and it is also known that depletion of HIRA might increase H3.3 pool leading to chromatin rearrangement (Choi et al., 2024). We anticipate that wherever we visualize a relaxation of the chromatin, other factor including incorporation of H3.3 might be responsible for the chromatin state which demands further experimental evidence.

SETDB1 associated increase in H3K9me3 level upon HIRA downregulation point to another interesting finding. Recent report indicated that SETDB1 negatively regulate pro-leukemia genes and thereby suppress AML mediated by H3K9me3 (Ropa et al., 2020). Our result also indicates a similar outcome.

Although, we demonstrated the differential incorporation of H3.3 within the *SETDB1* promoter resulted in upregulation of SETDB1 expression thereby inducing the level of H3K9me3, but there are already studies conducted which provide ample evidence that HIRA itself can in fact regulate histone modification, as demonstrated in yeast (Anderson et al., 2009). So, we predict that other additional factors might be involved in this regulatory axis responsible for this chromatin status.

The major concern in leukemia is restricted differentiation and abnormal proliferation and our study indicated that upon downregulation of HIRA, expression of proliferation markers are significantly reduced (Majumder et al., 2019) with reduced chromatin accessibility and enhanced H3K9me3 incorporation (this study). Role of HIRA in erythroid differentiation has been well documented (Majumder et al., 2019; Murdaugh et al. 2021). But the differentiation to erythroid and megakaryocyte lineage might be differentially regulated. Further exploration into ChIP-sequencing of H3K9me3 within CML cell genome might clarify the understanding on this differential incorporation and thereby regulation.

Our investigation offers a thorough understanding of the function of HIRA in the regulation of the chromatin architecture of CML cells, which could be exploited to comprehend the progression of leukemia and to develop HIRA or the associated molecules in the HIRA-SETDB1-H3K9me3 axis as potential molecular targets in CML therapy.

## Acknowledgements

We thank Dr. Ananda Mukherjee for valuable inputs and suggestions for the work. The work is supported by Science & Engineering Research Board (SERB), Department of Science & Technology (#CRG/2021/002528), partially supported by, #CRG/2022/005438, ICMR (2021-9968/SCR/ADHOC-BMS), DBT (#BT/PR51143/MED/31/471/2023) and intramural fund from the institute aided by Department of Biotechnology. MBS received fellowship from UGC #191620091128.

## Authors’ contributions

MBS conducted majority of the experiments, collected and analyzed data, wrote manuscript. AD did experiments on cell culture, western blots and EdU study; DD conceived the idea, performed experiments, analyzed data, arranged for resources and space, co-wrote the manuscript with MBS and with permission from other authors submitted to the journal.

## Data Availability Statement

The datasets generated during and/or analysed during the current study are available from the corresponding author on reasonable request.

## Abbreviations

AML: Acute myeloid leukemia
ATAC-seq: Assay for transposase-accessible chromatin with sequencing
ATRX: Alpha-thalassemia/mental retardation, X-linked
BCL2: B-cell leukemia/lymphoma 2 protein.
CDH1: Cadherin −1
CBBPA: CCAAT Enhancer Binding Protein Alpha
CEBPE: CCAAT enhancer binding protein epsilon
CML: Chronic myeloid leukemia
CXCL8: C-X-C motif chemokine ligand 8
DAXX: Death domain associated protein
EREG: Epiregulin
FLIM-FRET: Fluorescence lifetime imaging microscopy - Forster resonance energy transfer
FRAP: Fluorescence recovery after photobleaching
GFP: Green Fluorescent protein
GLI2: GLI family zinc finger 2
H2CA8: H2A clustered histone 8
HIRA: Histone cell cycle regulator A
KLF1: Erythroid Krueppel-Like Transcription Factor
MRTFA: Myocardin Related Transcription Factor A
PCNA: Proliferating Cell Nuclear Antigen
PRAME: PRAME nuclear receptor transcriptional regulator
SETDB1: SET domain bifurcated histone lysine methyltransferase 1
TFRC: Transferrin Receptor
XRN2: 5’-3’-exoribonuclease 2

