## Supplementary material for "HIRA-SETDB1-H3K9me3 axis regulate chromatin architecture in chronic myeloid leukemia cells": Figure S1, Figure S2

**Supplementary Information**

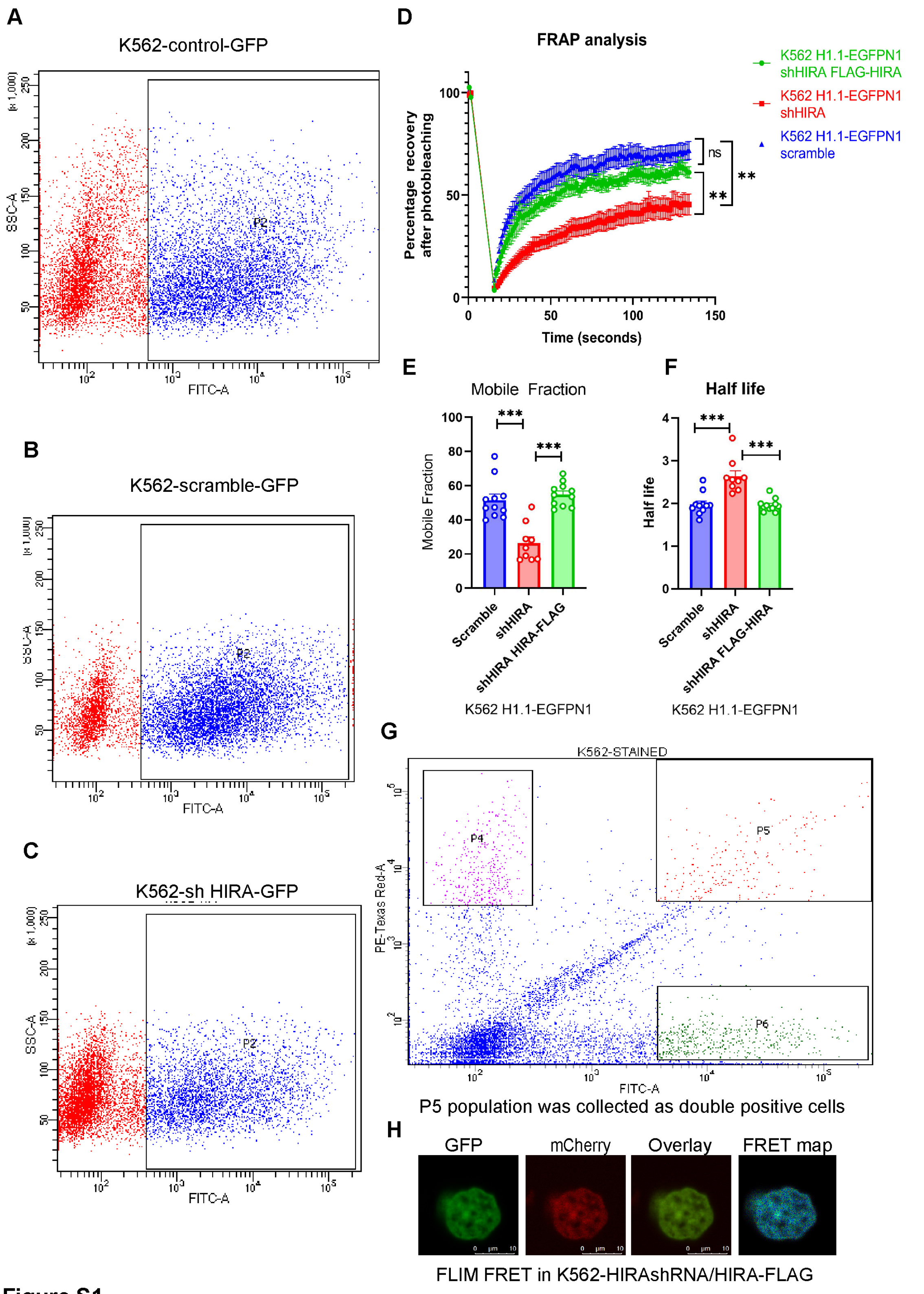


**Figure S1**

**Supplementary Figure S1.** A, B, C. FACS of H1.1-EGFP positive control, scramble-shRNA and HIRA-shRNA expressing K562 cells. P2 represent the sorted population used for FRAP analysis. D. FRAP analysis was conducted upon rescue of HIRA-shRNA with expression of HIRA-FLAG in the same cells. Fluorescence recovery after photobleacing demonstrates a similar pattern in rescued cells and scramble-shRNA expressing cells, while significant difference of the recovery pattern with HIRA-shRNA expressing cells. E, F. Bar graphs demonstrate the mobile fraction and half-time for the recovery of GFP in rescued cells in comparison to scramble and HIRA-shRNA expressing cells. Two-tailed unpaired T-test was performed for statistical analysis. ****p<0.0001. G. FACS for the H2B-mCherry/EGFP expressing K562 cells. P5 population was used for further experiments. H. FRET efficiency upon rescue of HIRA expression by HIRA-Flag in HIRA-shRNA K562 cells. Map shows similar efficiency map as in scramble-shRNA expressing K562 cells.

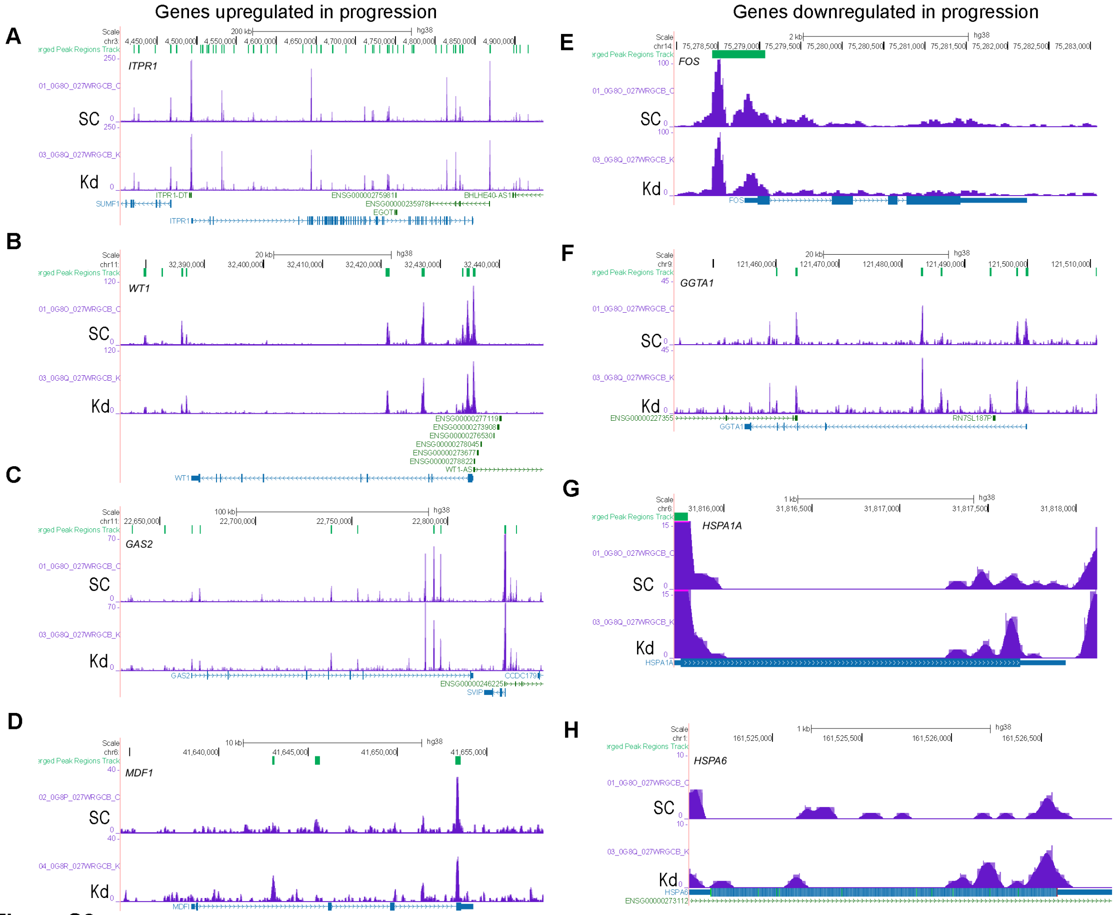


**Figure S2**

**Supplementary Figure S2.** A-D. ATAC sequencing for the chromatin accessibility within genomic loci upregulated during CML progression. E-H. ATAC sequencing for the chromatin accessibility within genomic loci downregulated during CML progression

**Supplementary video 1 and 2.** Z-stack image for the expression of H1.1 EGFP in K562 cells expressing scramble-shRNA and *HIRA*-shRNA respectively.

**Supplementary Tables**

**Table S1. Protein-protein interaction network.**

**Table S2. Cloning Primers**

| | |
|---|---|
| H1.1_EGFPN1 F | CCAAGCTTACCGCCATGTCTGAAACAGTGCCTCCCGC |
| H1.1-EGFPN1_R | CGGGATCCCCCTTTTTCTTGGGTGCCGC |
| H2B-EGFPC1_F | CCCTCGAGATGCCTGAACCCTCTAAGT |
| H2B_EGFPC1_R | GCGTCGACAGAGCTAGTGTACTTGG |
| HIRA_FLAG_F | CTCGAGATGAAGCTCCTGAAGCCGAC |
| HIRA_FLAG_R | GAATTCCTACTTGTCCCTCAGGATGTC |
| SETDB1_pEGFPC1_F | CGCTCGAGGTATGCCAACTTTGTACAAA |
| SETDB1_pEGFPC1_R | CGGTCGACTCAGAGATTTTGAGACAC |

**Table S3. ShRNA used in this study**

| | |
|---|---|
| shRNA_SETDB1_F | CCGGGCTCAGATGATAACTTCTGTACTCGAGTACAGAAGTTATCATCTG AGCTTTTTG |
| shRNA_SETDB1_R | AATTCAAAAAGCTCAGATGATAACTTCTGTACTCGAGTACAGAAGTTATC ATCTGAGC |
| shRNA_HIRA_F | CCGGCTCTATCCTCCGGAATCATTCCTCGAGGAATGATTCCGGAGGAT AGAGTTTTTG |
| shRNA_HIRA_R | AATTCAAAAACTCTATCCTCCGGAATCATTCCTCGAGGAATGATTCCGG AGGATAGAG |

**Table S4. ChIP primers used in this study**

| Gene name | Forward | Reverse |
|---|---|---|
| ITGA2B_promoter | ACAAGCGTTACTGTGAAGCG | GGGCCAGGAGACCTAAGAAATAA |
| SETDB1 promoter | CTGCCAGTCTCTTCTCACGT | CAGAGGCGACGAAAACTAAGG |
| TFRC | GGGGCCAGGCTATAAACCG | GATATCCCGACGCTCTGAGG |

**Table S5. Primary Antibody Used in this study**

| Primary antibody | Catalog number | Company | Dilutions |
|---|---|---|---|
| HIRA | 04-1488 | Merck Millipore | 1-1000 (WB) |
| GFP | 2956S | Cell Signaling Technology | 1-1000 |
| Ki67 | MA5-14520 | Thermo-Fisher Scientific | 1:2000 |
| H3 | 9715S | Cell Signaling Technology | 1-2000 |
| PCNA | SC7907 | Santacruz | 1:3000 |

| | | | |
|---|---|---|---|
| H3K9me3 | AB8898 | Abcam | 1:2000 (WB)<br>1:200 (IF) |
| H3K27me3 | AB6002 | Abcam | 1:2000 |
| H3K4me1 | | Abcam | 1:2000 |
| H3K4me2 | AB32356 | Abcam | 1:2000 |
| H3K4me3 | AB8580 | Abcam | 1:2000 |
| H3K36me3 | AB9050 | Abcam | 1:2000 |
| H3K27ac | AB4729 | Abcam | 1:2000 |
| H3K9ac | AB10812 | Abcam | 1:2000 |
| GAPDH | G9545 | Sigma | 1:10000 |
| SETDB1 | GTX115305 | Genetex | 1:3000 |
| SETDB1 | 11231—1-AP | Proteintech | 1:1000 |
| BCR-ABL | AB187831 | Abcam | 1:2000 |

**Secondary antibodies used in the study**

| Antibody | Company | Catalog number | Dilution |
|---|---|---|---|
| Goat anti mouse IgG-HRP | Santacruz | SC2005 | 1-2000 (WB) |
| Goat anti-rabit alexa-fluor 488 | Invitrogen | A11008 | 1-200 (IF) |
| Goat-anti-mouse alexa fluor 568 | Invitrogen | A11004 | 1-200 (IF) |

**Table S6. qRT-PCR primers used in this study**

| Gene name | Forward primer | Reverse primer |
|---|---|---|
| GAPDH | CACCAGGGCTGCTTTTAACTCTGGTA | CCTTGACGGTGCCATGGAATTTGC |
| KDM4B | AGACGTATGATGACATCGACGA | CGTAGATCGGGGAGACAAAGG |
| KDM4A | GAAGCCACGAGCATCCTATGA | GCGGAACTCTCGAACAGTCA |
| KDM4C | CGAGGTGGAAAGTCCTCTGAA | GGGCTCCTTTAGACTCCATGTAT |
| KDM4D | ATCGCCATTTGGCAAACAGTA | GGGTGTATTGACGCCTTCTATGA |
| KDM4E | AATCACGGCTTCAACTGCG | CATAACTCTCGGGTTGCACAA |
| SUV39H2 | TACTCGTCTTCCCCGAATAGC | GGCTGTGGTCAATAGAATCTGAA |
| SUV39H1 | CATCTGGGACGCATCACTGTA | TCACCAACACGGTACTCATTG |
| SETDB1 | TAAGACTTGGCACAAAGGCAC | TCCCCGACAGTAGACTCTTTC |
